# Intermingled representation of oral cavity in mouse trigeminal ganglion

**DOI:** 10.1101/2025.06.03.657656

**Authors:** Ashley Matunis, Ryotaro Iwamoto, Emma Stacy, Kenta Abe, Shunki Tamura, Yuki Kambe, Takahide Itokazu, Takatoshi Hikida, Tatsuo Sato, Takashi Sato

## Abstract

Somatotopy serves as a fundamental principle underlying sensory information processing, traditionally emphasized in the study of the cerebral cortex. However, little effort has been directed towards unraveling the spatial organization characterizing the earlier stages of sensory pathways. In this study, we developed a novel methodology to visualize individual neurons within the trigeminal ganglion—a crucial cluster of cell bodies of sensory neurons innervating the face. Our investigations revealed a reliable sensory response to stimulation of the lower teeth or lip within this ganglion. The responsive neurons were confined to a specific portion of the trigeminal ganglion, consistent with innervation of the lower oral cavity by the mandibular nerve. Contrary to our expectations, we did not observe a discernible map delineating specific regions of the oral cavity. Instead, the spatial representation of the teeth and lips exhibited unexpected intermingling. These findings challenge conventional understandings rooted in cortical maps and suggest that such conceptual frameworks may not be applicable to earlier stages of sensory pathways for the oral cavity. Our study sheds light on the complex spatial organization of sensory processing in the trigeminal system, highlighting the need for further research to elucidate the underlying mechanisms and implications for sensory perception and clinical interventions.

**Impact statement:** *In vivo* calcium imaging revealed that neurons responding to tooth stimulation are sparsely distributed in the trigeminal ganglion, intermixed with neurons responding to lip stimulation.

## Introduction

The sensations of teeth play a crucial role in controlling oral behaviors such as biting and chewing. For example, accurate perception of the load on the teeth is essential for the incisors to function as cutting tools, effectively splitting food morsels into smaller pieces with controlled force. The sensory signals applied to the teeth are primarily detected by mechanoreceptors in the periodontal ligaments between the root of the tooth and the alveolar bone ^1^, although they may also involve those in the tooth pulp ^2-4^. Consistently, studies have shown that humans lacking periodontal mechanoreceptors or under local anesthesia exhibit deficits in precise manipulative actions involving the application of low forces by the jaw ^5,6^.

The trigeminal ganglion (TG) contains the first group of neurons that receive sensory information from the face, including the oral cavity, and convey it to the brainstem ^7-10^. The TG constitutes the point of divergence for the three branches of the trigeminal nerve. The ophthalmic branch (V1) innervates the upper 1/3 of the facial region, from the forehead to the nose. The maxillary branch (V2) processes sensory information from the middle 1/3 of the facial region, including the upper jaw. The mandibular branch (V3) is responsible for sensory processing of the lower 1/3 of the facial region and further branches into smaller nerves, including the inferior alveolar nerve, which innervates the lower jaw, periodontal ligaments, and lower lip. Sensory information gathered from each part of the face/oral area, including signals from periodontal mechanoreceptors, is conveyed via neurons in the TG to the trigeminal nucleus in the brainstem, then to the thalamus, and finally to the somatosensory cortex.

Topographical organization is a fundamental guiding principle for sensory information processing ^11^. Within the somatosensory system, the somatosensory cortex contains topographic maps of the body surface, often depicted as the “homunculus” ^8,12-15^. Studies have demonstrated that distinct portions of the somatosensory cortex encode different parts of the oral cavity, such as the teeth, lips, and tongue ^16-20^. However, little is known about how such parts are spatially represented earlier in the somatosensory pathway, with only a limited number of studies tackled this question with extracellular unit recording ^21,22^ and anatomical tracing ^23^, primarily on the vibrissa system. In the present study, we employed *in vivo* calcium imaging ^12,13,24-28^ to investigate the spatial organization of the teeth and lips in the TG. We report that, in contrast to the cerebral cortex, the organization of the teeth and lips in the TG is spatially intermingled.

## Methods Animals

All experimental procedures were approved by the Medical University of South Carolina and Kagoshima University and conducted in conformity with the Institutional Animal Care and Use Committee at each institute. The experiments are in accordance with ARRIVE guidelines (https://arriveguidelines.org). Thy1-GCaMP6f mice (GP5.17, Jax #025393, both male and female, crossed with wild-type C57BL/6) were used in this study. Mice of both sexes, aged >8 weeks were included. The mice were maintained in group housing (up to five mice per cage) and experiments were performed during the dark period of a 12-h light/12-h dark cycle.

### Surgery

Mice were head-fixed under anesthesia (1%–5% isoflurane). A headpost was attached to the skull with glue and dental cement for stabilization during imaging. A craniotomy was performed above the left cerebral cortex, and cortical and subcortical tissues were removed via suction until the TG became visible. Then, a 3 × 3 × 8 mm glass cuboid was inserted above the TG. Small pieces of SURGIFOAM™ were placed around the cuboid to prevent bleeding into the imaging surface and to stabilize the glass cuboid. Glue and dental cement were applied around the glass cuboid to further secure it in place. Once the glass cuboid was secured, the animal was injected with an anesthetic (0.2 mL ketamine/xylazine cocktail) and placed under a fluorescence microscope (MVX, Olympus) for one-photon imaging. After imaging, the animals were deeply anesthetized with isoflurane and euthanized via transcardiac perfusion with 4% paraformaldehyde (PFA) in phosphate-buffered saline (PBS). The brain was then removed.

### Histology and Immunostaining

The perfused brain was placed in 30% sucrose solution after overnight fixation in 4% PFA. NeuN and GCaMP was immunostained using standard procedures ^26,29^ (Fig. 1b). Slices (thickness, 20 µm) were cut using a cryostat and blocked in carrier solution (5% bovine serum albumin, Sigma-Aldrich; 0.3% Triton X-100, Sigma-Aldrich; in 0.1 M PBS) for 1 h at room temperature on a shaker. For GCaMP staining, slices were incubated with anti-GFP primary rabbit antibody (1:1000 in carrier solution; Cat. #A11122, ThermoFisher) for 18 h at 4 °C on a shaker. For NeuN staining, slices were incubated with anti-NeuN chicken antibody (1:500 in carrier solution; ABN91, Millipore). After three rinses with 0.1 M PBS for 30 min, sections were incubated with either Alexa-Fluor-488-conjugated anti-rabbit secondary antibody (A21206, ThermoFisher, Lot# 2668665; 1:500 in carrier solution) or Alexa-Fluor-568-conjugated anti-chicken secondary antibody (A11041, ThermoFisher, Lot#2674372) for 1 h at room temperature on a shaker. After a few additional rinses for 30 min in 0.1 M PBS, slices were mounted on slide glasses for imaging (Fluorescent mounting medium, S302380, Dako).

### Stimulation

Sensory stimulation was applied to the facial (labial) surface of the left mandibular incisor and the outer surface of the lower lip in the diastema area. These two sites were chosen because they can be accessed without providing sensory stimulation to other parts of the oral cavity. We administered 15 stimulations in 3 s (200 ms intervals), with a baseline period of 3 s before and after the trial. The strength of the simulation is 20∼40 mN. This process was repeated for 10 trials per stimulation site.

### Imaging

The imaging experiments were conducted under a fluorescence microscope (MVX-10, Olympus). The excitation light was provided by a blue LED (SOLIS-470, Thorlabs), and the emission light was detected by a monochromatic camera (Thorlabs). The microscope was equipped with a 1x objective lens (MV PLAPO 1x/0.25 NA). The filter cube contained an excitation bandpass filter of 470/40 nm, a dichroic filter of 495 nm, and an emission bandpass filter of 525/50 nm. The sampling rate of the camera was 20–30 frames per second for an image size of 512 pixel x 512 pixel.

### Data processing

The imaging data were analyzed by a custom-written MATLAB program, as previously described ^24-27^. Imaging data were processed for motion correction and registration. Cells were detected for region-of-interest (ROI) drawing using Suite2p ^30^ and a custom-made MATLAB program ^26^. A fluorescence trace was generated for each ROI and then normalized by the baseline fluorescence (1 s period before the sensory stimulus onset) to produce a ΔF/F trace.

For each neuron, the sensory response of each trial was defined as the mean ΔF/F for the last 1 s of the sensory stimulation. The responses to the tooth and the lip were analyzed separately. If the sensory response over the 10 trials was significantly larger than 0 (*t*-test, *p* < 0.05), the response amplitude of the neuron was defined as the mean sensory response; otherwise, the response amplitude was defined as 0. For each neuron, the selectivity index was calculated as (response amplitude to tooth - response amplitude to lip) / (response amplitude to tooth + response amplitude to lip). We determined the center of gravity of the sensory response by calculating the mean of the cell location weighted by the response amplitude. If the response amplitude of a given neuron to the tooth or the lip was more than 5 times that to the other, the neuron was considered selective (i.e., selectivity index < -0.66 or selectivity index > 0.66).

The size of the neuron was determined as follows. First, the mean DF/F map was calculated for each individual pixel. Then, the ΔF/F of a 61 × 61 pixel area centered on an individual cell was fitted to the following two-dimensional Gaussian function.

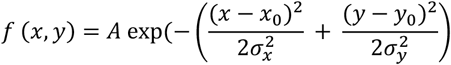

This fitting was accomplished using the MATLAB function lsqcurvefit.

## Statistical Analysis

Data are described as the mean ± s.e.m. unless otherwise noted. The significance of the response amplitude was determined by t-tests using MATLAB. All other statistics were based on non-parametric tests (Mann-Whitney test, Wilcoxon signed-rank test). Significance levels of data are denoted as * *p* < 0.05, ** *p* < 0.01, and *** *p* < 0.001. *p* > 0.05 was considered insignificant and is denoted as n.s.

## Results

To investigate the spatial organization of neurons encoding mechanical stimulation of the tooth, we employed *in vivo* calcium imaging, a method widely utilized in studying the topographical organization of sensory cortices ^12,13,31-33^ (Figure 1a). Several studies have achieved optical access to the TG by removing all of the superior structures and exposing it, often followed by a bath application of HEPES buffer, similar to *in vitro* preparations ^34-37^. In the current study, we modified a procedure that has been employed for two-photon imaging of the dorsal cerebral cortex ^38^. Specifically, we inserted an elongated glass cuboid (3 mm x 3 mm x 8 mm) directly above the TG after removing the overlying tissues (Figure 1c). This approach resulted in a broad and clear view of the dorsal surface of the TG (Figure 1d). Similar to numerous studies employing *in vivo* imaging of the cerebral cortex, our approach preserved tissue health within the cranial cavity, eliminating the need for buffer application during imaging sessions. However, a drawback of this approach is the long distance (> 8 mm) from the cover glass surface to the imaging site. As this length exceeds the working distance of our objective lens (CFI75, Nikon, working distance of 3 mm), conducting two-photon imaging was not feasible. Therefore, in the current experiment, we employed wide-field fluorescence imaging. We note that the use of an objective lens with a longer working distance would enable the visualization of neurons in the TG with a two-photon microscope.

**Figure 1.**
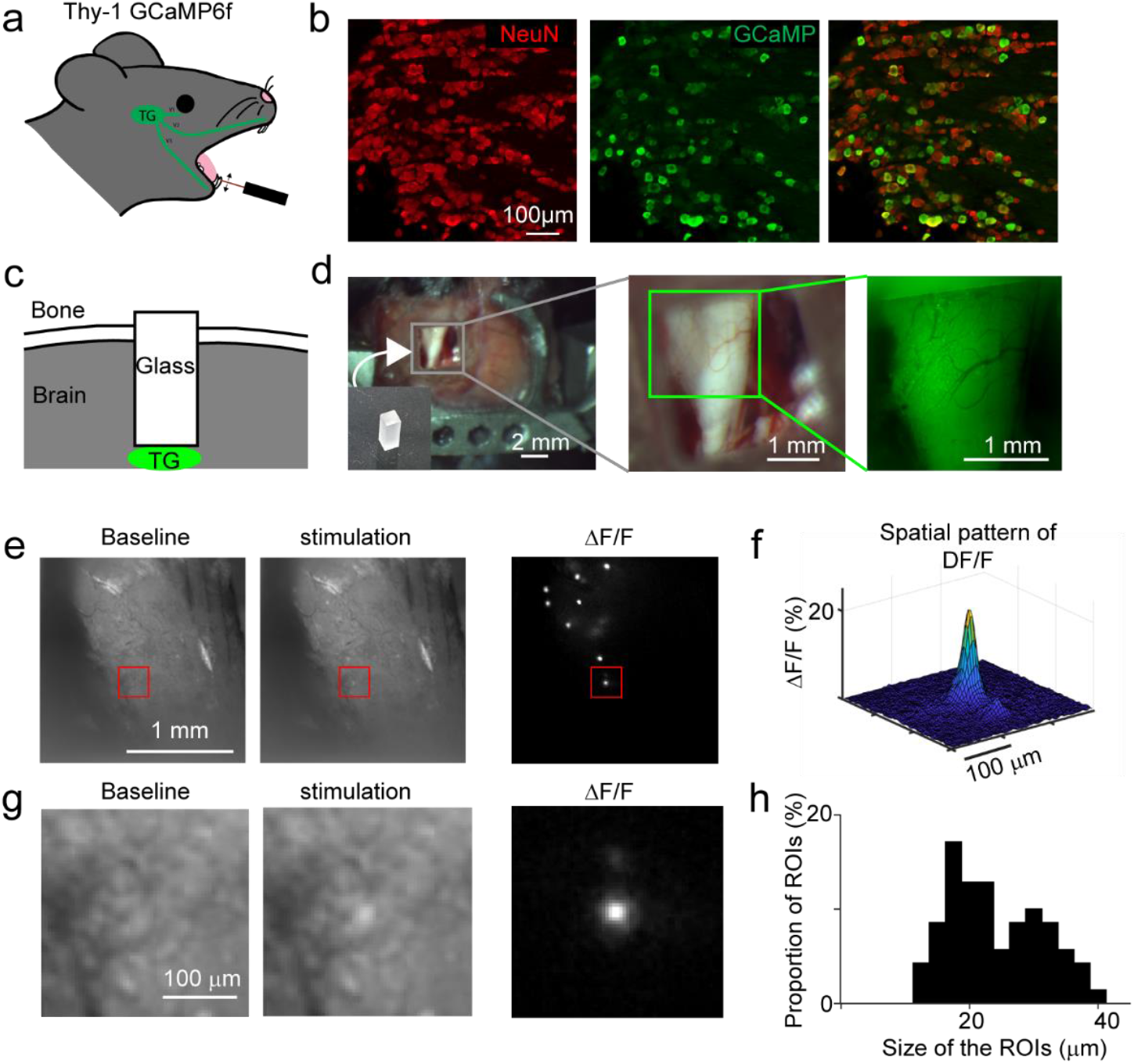
*In vivo* calcium imaging from the TG. (a) The distal processes of neurons of the TG innervate the face, including the oral cavity. Thus, stimulation of the tooth will activate the neurons in the TG. (b) Immunostaining for NeuN (red, left) and GCaMP (green, middle). The overlaid image is shown on the right.(c). Schematic view of our imaging approach. A 3 mm x 3 mm x 8 mm glass cuboid was placed through the brain above the TG (d) Surgical approach to the TG. The dorsal view of the surgical/imaging site (left). The TG was visible using both our surgical microscope (middle) and fluorescence microscope (right). (e) Examples of fluorescence images of the TG. Left, average image during the baseline period before sensory stimulation. Middle, average image during sensory stimulation. Right, spatial map of ΔF/F during sensory stimulation of the oral cavity. Increased signals were observed in some cells. (f) Three-dimensional representation of ΔF/F shown in panel e (g) High zoom images of the TG shown in panel e. The location is indicated by a red square in panele. (h) Distribution of the diameter of the area exhibiting increased fluorescence. The estimation of the diameter was based on Gaussian fitting.

We conducted fluorescence imaging of GCaMP6f in the TG of Thy-1 GCaMP mice (GP5.17) ^39^. Immunostaining against NeuN, a neuron specific marker ^40^, confirmed that the expression of GCaMP was restricted to neurons in these mice (Figure 1b). We successfully visualized several neurons through the glass cuboid, and some of these neurons exhibited fluorescence changes following manual stimulation of the oral cavity (Figure 1e,g, Supplementary Movie). The size of the activated area was 22.7 ± 0.9 µm (N = 70 neurons from 4 mice) (Figure 1f,h). Using this *in vivo* calcium imaging strategy, we examined the representation of teeth in the TG and compared it with the representation of the lips. We applied mechanical stimulation to either the facial (labial) surface of the left mandibular incisor (the front tooth located on the jaw, adjacent to the midline of the face) or the outside of the lower lip in the diastema area (between the incisors and molars) 15 times in 3 s (Figure 2a, top left). These two structures provided easy access, minimizing the risk of inadvertent stimulation of surrounding regions of the oral cavity. In contrast, the molars at the back of the mouth were difficult to approach for precise sensory stimulation. We chose to stimulate 15 times because we were unable to detect reliable fluorescence changes when we applied only single stimulations, presumably due to the lower signal-to-noise ratio of GCaMP6f compared to more recent versions of GCaMP, such as GCaMP8 ^41^.

**Figure 2.**
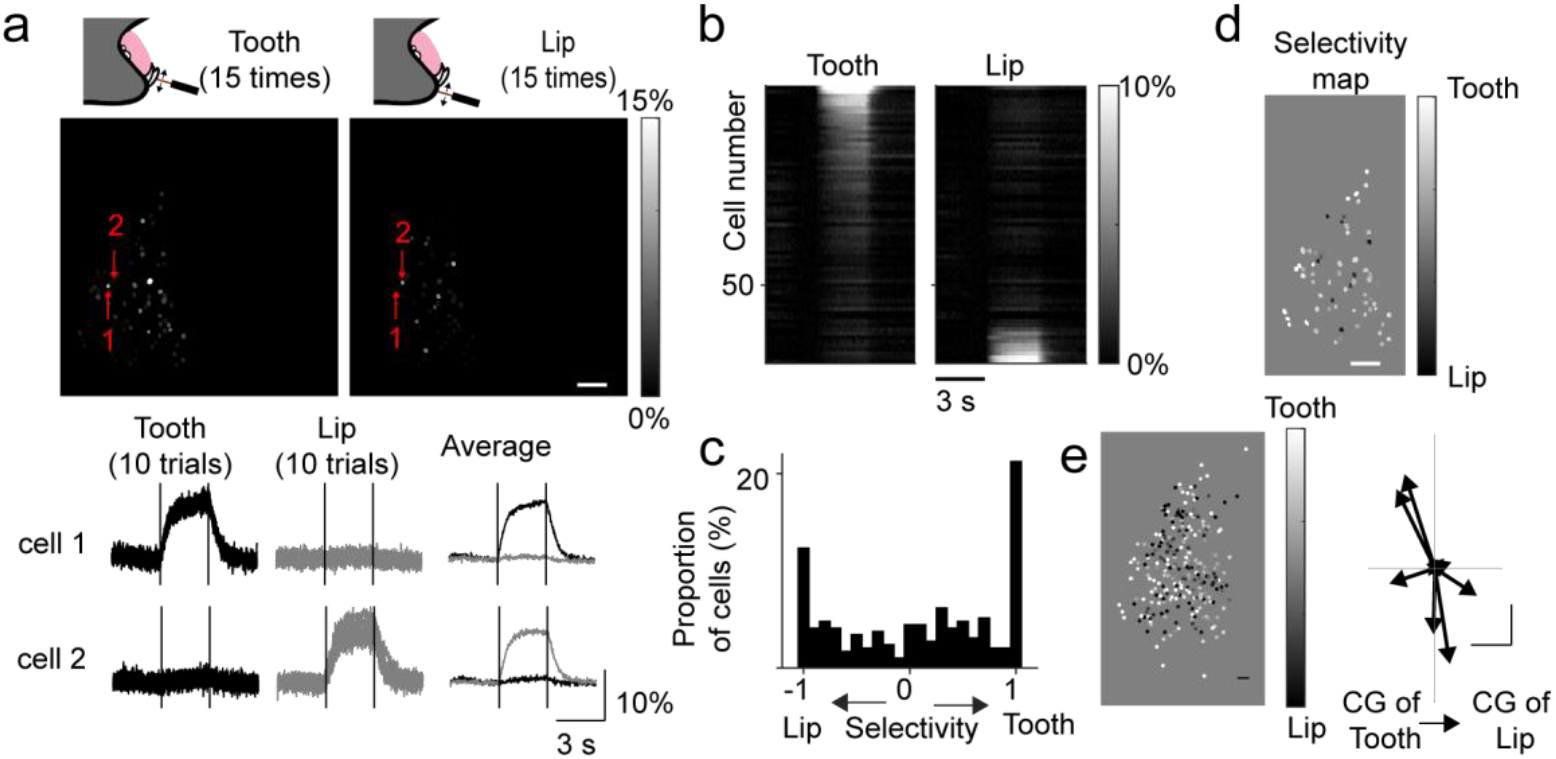
Spatial representation of the tooth and the lip in the TG. (a) Top, spatial maps of DF/F during tooth stimulation (left) and lip stimulation (right) from the same imaging site. Bottom, examples of fluorescent changes in two neurons indicated in the top panels. The left two columns show individual traces overlayed for 10 trials, and the right column shows the average traces. The locations of the two neurons are indicated. (b) ΔF/F traces for 69 neurons in panel a showing the responses to tooth stimulation (left) and lip stimulation (right). The cells were sorted based on the difference in activity during tooth and lip stimulation. (c) Distribution of the selectivity index from all imaging sessions (n = 292 from 9 mice). (d) Map of the selectivity index. Each circle corresponds to a cell, and the darkness of the cells indicates the selectivity index. White cells and black cells are intermingled. (e) Left, location of neurons overlaid from 9 experiments with the center of gravity of the tooth response placed at the center of the image (*n* = 292). There was no segregation between tooth-selective cells and lip-selective cells. Right, the relationship between the center of gravity for the tooth response and that for the lip response for 9 mice. There was no clear spatial relationship between them.

During sensory stimulation, the neurons exhibited increased fluorescence signals that decayed toward the baseline as the stimulation ended (Figure 2a, bottom left). Neurons responding to incisor stimulation and those activated by lower lip stimulation were predominantly localized at the entrance of the mandibular nerve ^35^, consistent with the innervation of the lower jaw, including these two structures, by the mandibular nerve of the TG. While the distribution of the active neurons was relatively sparse, the activity patterns of individual neurons were reproducible across multiple trials (Figure 2a, bottom left). For instance, in cell 1 of Figure 2a, the coefficient of variation (standard deviation/mean) for tooth responses was 0.044, and in cell 2 for lip responses, it was 0.164. Overall, the coefficient of variation of neurons exhibiting more than ΔF/F > 10% was 0.171 ± 0.036 (n = 16) for responses to incisor stimulation and 0.201 ± 0.042 (n = 39) for responses to lip stimulation. Notably, these coefficients of variation were substantially smaller than those reported in the cerebral cortex (e.g., refs ^42,43^).

We then compared the spatial patterns of sensory responses in the TG following stimulation of the incisor and the lower lip. Toward this goal, we first examined the response selectivity of individual neurons to incisor and lip stimulation. The majority of responsive neurons exhibited selectivity to either incisor or lip stimulation, with 53.4 % of neurons responding at least four times more to one stimulus than the other (*n* = 156/292) (Figure 2b,c). We classified these neurons as tooth-selective neurons and lip-selective neurons. However, a significant number of neurons responded to both the lower incisor and the lower lip (Figure 2c). Spatially, the tooth-selective neurons and lip-selective neurons did not form clusters; instead, they tended to be intermingled (Figure 2d,e). When we aligned all imaging sessions based on the center of gravity of tooth responses (Figure 2e), we found similar distances to tooth-selective cells and lip-selective cells (370.4 ± 24.0 µm, *n* = 84 vs. 343.3 ± 20.0 µm, *n* = 72, *p* = 0.448, Mann-Whitney U test). Furthermore, no consistent spatial relationship emerged between the center of gravity for tooth responses and that for lip responses (in the horizontal direction, *p* = 0.82, in the vertical direction *p* = 0.91, *n* = 9, Wilcoxon signed-rank test) (Figure 2e, right). These findings collectively indicate a lack of clear topological organization of tooth and lip representations at the level of the TG.

## Discussion

In this study, we introduced a novel approach involving the implantation of an elongated glass cuboid to visualize the sensory representation of neurons in the TG. This innovative method enabled us to explore the spatial organization of tooth-responsive neurons and revealed an unexpected finding: the spatial intermingling of neurons responsive to incisor and lip stimulation. The weak sensory stimulation used in our experiment activates primarily low threshold mechanoreceptive afferents ^44^, and the sensory map for other cell-types ^45^ may exhibit distinct spatial patterns. Nonetheless, our findings represent a crucial initial step toward understanding the processing of sensory information related to teeth in the brain, presenting a new avenue for investigating the neural circuitry underlying toothaches—a pervasive issue affecting a vast portion of the global population.

Our approach offers several advantages over traditional methods used to visualize the TG. Unlike previous techniques involving the removal of structures above the TG and the use of HEPES buffer ^34-37^, our method maintains the TG within a contained area in the cranial cavity, eliminating the need for buffer supplementation during experiments. Additionally, the direct contact of the glass cuboid with the TG enhances imaging site stability against mouse movement induced by breathing and heartbeat, akin to approaches used in visualizing the cerebral cortex ^26,28,38,46^. Furthermore, our approach holds promise for long-term imaging across multiple days, despite the removal of a significant portion of the brain.

Despite its advantages, our approach has limitations that warrant consideration. First, due to the length of glass cuboid (8 mm), the approach was not compatible with *in vivo* two-photon imaging because our current objective lens has a working distance of 3 mm. This limitation restricts the signals we obtain to those originating primarily from the surface of the TG, impeding our ability to detect spatial organization in the dorsal-ventral direction. For instance, a recent study showed that tongue-innervating neurons are located on the ventral side of the TG ^47^, and we indeed could find minimum neural responses to the tongue stimulation in our experiments. To overcome this limitation along the dorsal-ventral axis, future studies will require *in vivo* two-photon imaging with an objective lens with a working distance of ∼8 mm. Second, filling the space with glass that has a high refractive index could cause spherical aberration. Although such aberration did not hinder our single-cell resolution imaging, future studies aiming to investigate smaller structures may need to employ an objective lens with correction collars to compensate for the refractive index mismatch.

We discovered that some neurons responded to both the tooth and the lip, although for most neurons, the response was substantially larger for one stimulus. Previous studies have reported that neurons in the trigeminal nucleus, the area postsynaptic to the neurons we imaged in the current study, also respond to multiple structures in the oral cavity ^48^. There are two possibilities: first, our mechanical stimulation was transmitted to a different portion of the oral cavity; second, individual neurons in the TG innervate both the lip and the incisor. Although our stimulation protocol was controlled to produce responses in only a subset of neurons, we cannot entirely exclude the first possibility. Determining how the receptive field structures of the oral cavity are formed in the TG will require an approach that employs single-cell labeling.

Earlier studies on vibrissae system have proposed a rough organization of whiskers in the TG ^21,22^, but due to the technical limitation of extracellular unit recording, the precise spatial patterns of sensory responses was not known. Our studies using *in vivo* calcium imaging provide strong evidence for an intermingled representation of the oral cavity in the TG. Our results on the TG contrasts with studies on the somatosensory cortex, which reported distinct representations of the lip and tooth as part of the “homunculus” in this brain region ^16-20^. Future investigations will explore how spatially distributed neuronal activity in the TG converges onto specific cortical regions via the thalamus, providing insights into higher-order sensory processing mechanisms.

In summary, our study introduces a novel approach to investigating the spatial organization of neuronal responses in the TG. The unexpected finding of spatial intermingling of tooth and lip representations challenges existing paradigms of somatosensory processing and sets the stage for further exploration of the complex neural circuitry underlying oral sensory perception.

## Author contributions

A.M., R.I, H.T., T.K.S., and T.R.S. conceived the project and designed the experiments. T.R.S. and T.K.S. prepared the analysis codes. A.M., R.I., E.S., and K.A. carried out the experiments. A.M., R.I., E.S., K.A., Y.K., T.H., T.K.S., and T.R.S analysed the data and wrote the manuscript.

## Resource Availability Lead Contact

Further information and requests for reagents may be directed to the Lead Contact, Takashi R. Sato.

## Materials Availability

These studies did not generate unique reagents.

## Data and Code Availability

All data and code used for figure generation in this study have been deposited in GitHub. http://github.com/pharmedku/trigeminal

## Acknowledgements

This work was supported by grants from the Brain and Behavior Research Foundation (Young Investigator Grant, 29268), National Institute on Drug Abuse (COCA pilot grant, P50 DA046373), National Institute of Aging (R03 AG070517), National Institute of Neurological Disorders and Stroke (R21 NS125571, R01 NS131549), and NIH COBRE in Neurodevelopment and its Disorders (P20 GM148302) to T.R.S.; AMED (JP21wm0425010, JP21gm1510006) and KAKENHI (JP22H02944, JP23K18163), the Collaborative Research Program of Institute for Protein Research, Osaka University, ICR-24-03 to T.H.; JST PRESTO (JPMJPR1883) and KAKENHI (20K23378) to T.K.S.

## References

1 Jacobs, R. & van Steenberghe, D. Role of periodontal ligament receptors in the tactile function of teeth: a review. J Periodontal Res 29, 153–167 (1994). 10.1111/j.1600-0765.1994.tb01208.x

2 Dong, W. K. & Chudler, E. H. Origins of tooth pulp-evoked far-field and early near-field potentials in the cat. J Neurophysiol 51, 859–889 (1984). 10.1152/jn.1984.51.5.859

3 Dong, W. K., Shiwaku, T., Kawakami, Y. & Chudler, E. H. Static and dynamic responses of periodontal ligament mechanoreceptors and intradental mechanoreceptors. J Neurophysiol 69, 1567–1582 (1993). 10.1152/jn.1993.69.5.1567

4 Dong, W. K., Chudler, E. H. & Martin, R. F. Physiological properties of intradental mechanoreceptors. Brain Res 334, 389–395 (1985). 10.1016/0006-8993(85)90239-2

5 Trulsson, M. & Johansson, R. S. Forces applied by the incisors and roles of periodontal afferents during food-holding and -biting tasks. Exp Brain Res 107, 486–496 (1996). 10.1007/BF00230428

6 Trulsson, M. & Gunne, H. S. Food-holding and -biting behavior in human subjects lacking periodontal receptors. J Dent Res 77, 574–582 (1998). 10.1177/00220345980770041001

7 Purkart, L. et al. Trigeminal ganglion and sensory nerves suggest tactile specialization of elephants. Curr Biol 32, 904–910.e903 (2022). 10.1016/j.cub.2021.12.051

8 Petersen, C. C. The functional organization of the barrel cortex. Neuron 56, 339–355 (2007). 10.1016/j.neuron.2007.09.017

9 Stüttgen, M. C., Kullmann, S. & Schwarz, C. Responses of rat trigeminal ganglion neurons to longitudinal whisker stimulation. J Neurophysiol 100, 1879–1884 (2008). 10.1152/jn.90511.2008

10 Severson, K. S., Xu, D., Yang, H. & O’Connor, D. H. Coding of whisker motion across the mouse face. Elife 8 (2019). 10.7554/eLife.41535

11 Kaas, J. H. Topographic maps are fundamental to sensory processing. Brain Res Bull 44, 107–112 (1997). 10.1016/s0361-9230(97)00094-4

12 Sato, T. R. & Svoboda, K. The functional properties of barrel cortex neurons projecting to the primary motor cortex. J Neurosci 30, 4256–4260 (2010). 10.1523/JNEUROSCI.3774-09.2010

13 Sato, T. R., Gray, N. W., Mainen, Z. F. & Svoboda, K. The functional microarchitecture of the mouse barrel cortex. PLoS Biol 5, e189 (2007). 10.1371/journal.pbio.0050189

14 Penfield, W. & Rasmussen, T. The cerebral cortex of man; a clinical study of localization of function. (Macmillan, 1950).

15 Brecht, M. The Body Model Theory of Somatosensory Cortex. Neuron 94, 985–992 (2017). 10.1016/j.neuron.2017.05.018

16 Taira, K. The representation of the oral structures in the first somatosensory cortex of the cat. Brain Res 409, 41–51 (1987). 10.1016/0006-8993(87)90739-6

17 Cusick, C. G., Wall, J. T. & Kaas, J. H. Representations of the face, teeth and oral cavity in areas 3b and 1 of somatosensory cortex in squirrel monkeys. Brain Res 370, 359–364 (1986). 10.1016/0006-8993(86)90494-4

18 Catania, K. C. & Remple, M. S. Somatosensory cortex dominated by the representation of teeth in the naked mole-rat brain. Proc Natl Acad Sci U S A 99, 5692–5697 (2002). 10.1073/pnas.072097999

19 Jain, N., Qi, H. X., Catania, K. C. & Kaas, J. H. Anatomic correlates of the face and oral cavity representations in the somatosensory cortical area 3b of monkeys. J Comp Neurol 429, 455–468 (2001). 10.1002/1096-9861(20010115)429:3<455::aid-cne7>3.0.co;2-f

20 Remple, M. S., Henry, E. C. & Catania, K. C. Organization of somatosensory cortex in the laboratory rat (Rattus norvegicus): Evidence for two lateral areas joined at the representation of the teeth. J Comp Neurol 467, 105–118 (2003). 10.1002/cne.10909

21 Leiser, S. C. & Moxon, K. A. Relationship between physiological response type (RA and SA) and vibrissal receptive field of neurons within the rat trigeminal ganglion. J Neurophysiol 95, 3129–3145 (2006). 10.1152/jn.00157.2005

22 Zucker, E. & Welker, W. I. Coding of somatic sensory input by vibrissae neurons in the rat’s trigeminal ganglion. Brain Res 12, 138–156 (1969). 10.1016/0006-8993(69)90061-4

23 Maklad, A., Fritzsch, B. & Hansen, L. A. Innervation of the maxillary vibrissae in mice as revealed by anterograde and retrograde tract tracing. Cell Tissue Res 315, 167–180 (2004). 10.1007/s00441-003-0816-z

24 Sato, T. R. et al. Interhemispherically dynamic representation of an eye movement-related activity in mouse frontal cortex. Elife 8 (2019). 10.7554/eLife.50855

25 Niethard, N. et al. Sleep-Stage-Specific Regulation of Cortical Excitation and Inhibition. Curr Biol 26, 2739–2749 (2016). 10.1016/j.cub.2016.08.035

26 Itokazu, T. et al. Streamlined sensory motor communication through cortical reciprocal connectivity in a visually guided eye movement task. Nat Commun 9, 338 (2018). 10.1038/s41467-017-02501-4

27 Hasegawa, M. et al. Selective Suppression of Local Circuits during Movement Preparation in the Mouse Motor Cortex. Cell Rep 18, 2676–2686 (2017). 10.1016/j.celrep.2017.02.043

28 Abe, K. et al. Enhanced Aversive Signals During Classical Conditioning in Dopamine Axons in Medial Prefrontal Cortex. bioRxiv (2023). 10.1101/2023.08.23.554475

29 Sato, T., Tokuyama, W., Miyashita, Y. & Okuno, H. Temporal and spatial dissociation of expression patterns between Zif268 and c-Fos in rat inferior olive during vestibular compensation. Neuroreport 8, 1891–1895 (1997). 10.1097/00001756-199705260-00020

30 Pachitariu M et al. Suite2p: beyond 10,000 neurons with standard two-photon microscopy. bioRxiv (2016).

31 Ohki, K., Chung, S., Ch’ng, Y. H., Kara, P. & Reid, R. C. Functional imaging with cellular resolution reveals precise micro-architecture in visual cortex. Nature 433, 597–603 (2005). 10.1038/nature03274

32 Rothschild, G., Nelken, I. & Mizrahi, A. Functional organization and population dynamics in the mouse primary auditory cortex. Nat Neurosci 13, 353–360 (2010). 10.1038/nn.2484

33 Bandyopadhyay, S., Shamma, S. A. & Kanold, P. O. Dichotomy of functional organization in the mouse auditory cortex. Nat Neurosci 13, 361–368 (2010). 10.1038/nn.2490

34 Ghitani, N. et al. Specialized Mechanosensory Nociceptors Mediating Rapid Responses to Hair Pull. Neuron 95, 944–954.e944 (2017). 10.1016/j.neuron.2017.07.024

35 Yarmolinsky, D. A. et al. Coding and Plasticity in the Mammalian Thermosensory System. Neuron 92, 1079–1092 (2016). 10.1016/j.neuron.2016.10.021

36 Hu, M., Doyle, A. D., Yamada, K. M. & Kulkarni, A. B. Visualization of trigeminal ganglion sensory neuronal signaling regulated by Cdk5. Cell Rep 38, 110458 (2022). 10.1016/j.celrep.2022.110458

37 Moayedi, Y. et al. The cellular basis of mechanosensation in mammalian tongue. Cell Rep 42, 112087 (2023). 10.1016/j.celrep.2023.112087

38 Komiyama, T. et al. Learning-related fine-scale specificity imaged in motor cortex circuits of behaving mice. Nature 464, 1182–1186 (2010). 10.1038/nature08897

39 Dana, H. et al. Thy1-GCaMP6 transgenic mice for neuronal population imaging in vivo. PLoS One 9, e108697 (2014). 10.1371/journal.pone.0108697

40 Mullen, R. J., Buck, C. R. & Smith, A. M. NeuN, a neuronal specific nuclear protein in vertebrates. Development 116, 201–211 (1992). 10.1242/dev.116.1.201

41 Zhang, Y. et al. Fast and sensitive GCaMP calcium indicators for imaging neural populations. Nature 615, 884–891 (2023). 10.1038/s41586-023-05828-9

42 Beaman, C. B., Eagleman, S. L. & Dragoi, V. Sensory coding accuracy and perceptual performance are improved during the desynchronized cortical state. Nat Commun 8, 1308 (2017). 10.1038/s41467-017-01030-4

43 Jones, L. M., Fontanini, A., Sadacca, B. F., Miller, P. & Katz, D. B. Natural stimuli evoke dynamic sequences of states in sensory cortical ensembles. Proc Natl Acad Sci U S A 104, 18772–18777 (2007). 10.1073/pnas.0705546104

44 Trulsson, M. Sensory-motor function of human periodontal mechanoreceptors. J Oral Rehabil 33, 262–273 (2006). 10.1111/j.1365-2842.2006.01629.x

45 Yang, L. et al. Human and mouse trigeminal ganglia cell atlas implicates multiple cell types in migraine. Neuron 110, 1806–1821.e1808 (2022). 10.1016/j.neuron.2022.03.003

46 Dombeck, D. A., Khabbaz, A. N., Collman, F., Adelman, T. L. & Tank, D. W. Imaging large-scale neural activity with cellular resolution in awake, mobile mice. Neuron 56, 43–57 (2007). 10.1016/j.neuron.2007.08.003

47 Zhang, L., Nagel, M., Olson, W. P., Chesler, A. T. & O’Connor, D. H. Trigeminal innervation and tactile responses in mouse tongue. Cell Rep 43, 114665 (2024). 10.1016/j.celrep.2024.114665

48 Sessle, B. J. & Greenwood, L. F. Inputs to trigeminal brain stem neurones from facial, oral, tooth pulp and pharyngolaryngeal tissues: I. Responses to innocuous and noxious stimuli. Brain Res 117, 211–226 (1976). 10.1016/0006-8993(76)90731-9

